# The global burden of trichiasis in 2016

**DOI:** 10.1101/348995

**Authors:** Rebecca M Flueckiger, Paul Courtright, Mariamo Abdala, Amza Abdou, Zaid Abdulnafea, Tawfik K Al-Khatib, Khaled Amer, Olga Nelson Amiel, Sossinou Awoussi, Ana Bakhtiari, Wilfried Batcho, Assumpta Lucienne Bella, Kamal Hashim Bennawi, Simon J Brooker, Brian K Chu, Michael Dejene, Djore Dezoumbe, Balgesa Elkheir Elshafie, Aba Ange Elvis, Djouma Nembot Fabrice, Fatma Juma Omar, Missamou François, Drabo François, Jambi Garap, Michael Gichangi, André Goepogui, Jaouad Hammou, Boubacar Kadri, George Kabona, Martin Kabore, Khumbo Kalua, Mathias Kamugisha, Biruck Kebede, Kaba Keita, Asad Aslam Khan, Genet Kiflu, Garae Mackline, Colin Macleod, Portia Manangazira, Michael P Masika, Marilia Massangaie, Taka Fira Mduluza, Nabicassa Meno, Nicholas Midzi, Abdallahi Ould Minnih, Sailesh Mishra, Caleb Mpyet, Nicholas Muraguri, Upendo Mwingira, Beido Nassirou, Jean Ndjemba, Cece Nieba, Jeremiah Ngondi, Nicholas Olobio, Alex Pavluck, Isaac Phiri, Rachel Pullan, Babar Qureshi, Boubacar Sarr, Do Seiha, Gloria Marina Serrano Chávez, Shekhar Sharma, Siphetthavong Sisaleumsak, Khamphoua Southisombath, Gretchen Stevens, Andeberhan Tesfazion Woldendrias, Lamine Traoré, Patrick Turyaguma, Rebecca Willis, Georges Yaya, Souleymane Yeo, Francisco Zambroni, Jialiang Zhao, Anthony W Solomon

## Abstract

**Background:** Trichiasis is present when one or more eyelashes touches the eye. Uncorrected, it can cause blindness. Accurate estimates of numbers affected, and their geographical distribution, help guide resource allocation.

**Methods:** We obtained district-level trichiasis prevalence estimates for 44 endemic and previously-endemic countries. We used (1) the most recent data for a district, if more than one estimate was available; (2) age- and sex-standardized corrections of historic estimates, where raw data were available; (3) historic estimates adjusted using a mean adjustment factor for districts where raw data were unavailable; and (4) expert assessment of available data for districts for which no prevalence estimates were available.

**Findings:** Internally age- and sex-standardized data represented 1,355 districts and contributed 662 thousand cases (95% confidence interval [CI] 324 thousand-1.1 million) to the global total. age- and sex-standardized district-level prevalence estimates differed from raw estimates by a mean factor of 0.45 (range 0.03-2.28). Previously non-standardized estimates for 398 districts, adjusted by ×0.45, contributed a further 411 thousand cases (95% CI 283-557 thousand). Eight countries retained previous estimates, contributing 848 thousand cases (95% CI 225 thousand-1.7 million). New expert assessments in 14 countries contributed 862 thousand cases (95% CI 228 thousand-1.7 million). The global trichiasis burden in 2016 was 2.8 million cases (95% CI 1.1-5.2 million).

**Interpretation:** The 2016 estimate is lower than previous estimates, probably due to more and better data; scale-up of trichiasis management services; and reductions in incidence due to lower active trachoma prevalence.

**Author Summary:** As an individual with trichiasis blinks, the eyelashes abrade the cornea, which can lead to corneal opacity and blindness. Through high quality surgery, which involves correcting the position of the in-turned eyelid, it is possible to reduce the number of people with trichiasis. An accurate estimate of the number of persons with trichiasis and their geographical distribution are needed in order to effectively align resources for surgery and other necessary services. We obtained district-level trichiasis prevalence estimates for 44 endemic and previously-endemic countries. We used the most recently available data and expert assessments to estimate the global burden of trichiasis. We estimated that in 2016 the global burden was 2.8 million cases (95% CI 1.1-5.2 million).

The 2016 estimate is lower than previous estimates, probably due to more and better data; scale-up of trichiasis management services; and reductions in incidence due to lower active trachoma prevalence.

## Background

Trachoma, a neglected tropical disease, is endemic or has recently been endemic in more than 50 countries[1]. It affects the most impoverished people of the world. Improved living standards are credited for the disappearance of trachoma from Europe and North America, but in many less developed countries, trachoma is still a public health problem, and contributes to the continued suffering and deepening of poverty of millions of people.

*Chlamydia trachomatis* is the causative organism. Repeated ocular chlamydial infection results in chronic inflammation, characterised by sub-epithelial follicles in the tarsal conjunctiva, which may be sufficiently large and numerous to meet the definition of trachomatous inflammation—follicular (TF), a key sign for assessing trachoma prevalence[2]. Over time, with repeated reinfection, scarring may develop; this scarring can eventually cause the eyelid to turn inwards in some people, resulting in eyelashes touching the globe. This is called trachomatous trichiasis (TT) and is very painful[3]. As an individual with trichiasis blinks, the eyelashes abrade the cornea, which can lead to corneal opacity and blindness[4].

The World Health Organization (WHO) endorses population based prevalence surveys (PBPSs) to guide decisions regarding implementation of interventions for trachoma elimination[5]. The typical survey design involves two-stage cluster sampling, which uses probability-proportional-to-size methodologies to select 20-30 clusters. The outputs are estimates of the prevalence of TF in children aged 1-9 years, and the prevalence of TT in adults aged 15 years and older[6].

Through high quality surgery, which involves correcting the position of the in-turned eyelid,[7] it is possible to reduce the number of people with TT. An accurate estimate of the number of persons with TT (the TT backlog) and their geographical distribution are needed in order to effectively align resources for surgery and other necessary services.

In 2009, Mariotti et al estimated the global TT backlog to be 8.2 million people[8]. This estimate was derived by summarizing a combination of published and unpublished information. First, a literature review was performed to identify published prevalence data. Where published data were not available, unpublished data were compiled from the Eleventh (2007) Meeting of the WHO Alliance for the Global Elimination of Trachoma by 2020[9]. Where information was still missing, unpublished reports were provided by health ministries. Finally, if none of these resources were available, data were extrapolated from a proxy country believed to have common epidemiological conditions and demographic structure.

There are many uncertainties inherent in the 2009 estimate[8]. First, where PBPS data were available, the results were not systematically standardized by age and sex. This is problematic because women and the elderly are more likely than men and younger adults, respectively, to (1) be at home at the time that a house-to-house survey team calls, and (2) have TT. Second, in countries for which data were available, survey coverage was generally far from complete. Though not an invariable rule, surveys to estimate the prevalence of neglected diseases have a tendency to be done first in areas of high prevalence; for the 2009 estimate, if any data were available for a particular country, the TT prevalence figure from it was applied across the yet-to-be-mapped suspected-endemic population. Third, the use of proxy countries is extremely subjective.

In 2012, WHO collected and collated provisional 2011 country reports, and published an updated figure for the global TT backlog. The total given was 7.3 million people, but the methodologies used in each country to generate national backlog figures were not described[10]; they are likely to have been highly heterogeneous.

From December 2012 to January 2016, the Global Trachoma Mapping Project (GTMP) sought to systematically complete the global trachoma map using standardized techniques for both collecting and analysing survey data[11]. GTMP measured trachoma prevalence using gold standard PBPSs conducted at district level. People of all ages living in selected households of 1,542 districts across 29 countries were examined for trachoma, resulting in the examination of 2.6 million people[12]. GTMP analyses included standardizing trichiasis prevalence estimates against national population pyramids in an attempt to partially account for demographic differences between those examined and the national averages.

As a result of the GTMP, there are now high quality PBPS data available for most suspected-endemic areas that were previously unsurveyed. The availability of these data has catalysed the current attempt to generate a new estimate of the global trichiasis backlog.

## Methodology

We updated previous global estimates using the best available data, according to the following hierarchy. First, where GTMP data[11] were available, we used the age- and sex-standardized trichiasis prevalence estimates for adults in each survey. In August 2014, GTMP added examination for the presence or absence of trachomatous scarring (TS)[2] for all eyes determined to have trichiasis; prior to this, and for all non-GTMP data, trichiasis prevalence estimates used must be considered to be estimates of “all-cause trichiasis” rather than “trachomatous trichiasis”.

Second, where PBPSs had been done without the support of the GTMP, we requested raw survey data from national health ministries, or appropriate government-designated agencies. Where those data were provided by 1 March 2016, we applied the same age- and sex-standardization as was used in the GTMP[11].

Third, where PBPSs had been done but raw data were not available, prevalence estimates (whether standardized for age and sex or not) were obtained from country programs. When data had not been standardized, estimates were multiplied by the mean adjustment factor for districts in which age- and sex-standardization was possible.

Fourth, if prevalence estimates were not available, we reviewed previous estimates through desk reviews and communications with country programs. If there was adequate evidence to revise the estimates, new estimates were used; otherwise, the 2009 estimates[8] were retained.

The population data used in this analysis were derived from the UN population division (UNdata)[13] and www.worldpop.org[14]. As trachoma is a disease primarily affecting rural populations[4], we used rural population estimates for this purpose. Rural population pyramids were obtained from UNdata. Microsoft Excel (2007) was used to organize the datasets into 5-year age bands stratified by sex for each country. The percentage of the population within each stratum was estimated from the bands. The district level populations were derived from www.worldpop.org raster files[14] using the zonal statistics tool in ArcGIS 10.3[15], where the sum raster value falling within each district boundary was taken as the district value. A sensitivity analysis was also performed, comparing the www.worldpop.org population estimates and district population estimates provided by national trachoma elimination programs. The mean ratio between the national program estimates and www.worldpop.org was 0.97. Because of this close correlation, and for the purposes of standardizing our methods, the www.worldpop.org data were used throughout this analysis.

The statistical software package R[16] was used to perform age- and sex-standardization. First, the crude prevalence was calculated for each cluster. Second, the prevalence was standardized for each cluster by weighting the proportion of each sex-specific five-year age band observed to have trichiasis by the proportion of the adults aged 15 years and older expected to have that age and sex in that district, if available, or (if not available), nation-wide. Third, the un-weighted arithmetic mean of the standardized cluster-level trichiasis proportions was taken as the district-level prevalence. Once the initial standardization was completed, bootstrapping was undertaken on the dataset for each individual district to derive 95% confidence intervals (CIs). For a district surveyed by examining individuals in *n* clusters, this involved bootstrap resampling (with replacement) of *n* clusters, over 10,000 replications. The R code is provided in the Appendix.

Where PBPSs had been done but raw data were not available, CIs were constructed by applying an approximation to the binomial distribution *CI* = *p* - (*ź* × *s. e.*) *to p* + (*ź* × *s. e.*), where p is proportion of positive cases in the sample, s.e. is 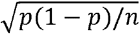, and *ź* is the standard normal distribution (95% CI = 1.96). Where the sample size was not available, a sample of 1000 was assumed.

Standardized prevalence estimates and the lower and upper 95% CI bounds were multiplied by the 15-years-and-older population for the relevant district (projected to 2016) to provide an estimate of the number of persons with trichiasis.

For expert backlog assessments, confidence intervals were constructed using the relative precisions of all datasets for which bootstrapped CIs could be determined. The range of relative upper bound and lower bound precisions were trimmed to exclude 10% of the greatest and least values. The means of the remainder were applied.

Upper (and lower) CI bounds around district-level estimates, regardless of the method used for calculation, were summed to generate country-level, region-level and global estimates of the upper (and lower) CI bounds.

## Role of the funding source and ethical review

The funder (WHO) of this project was involved in all aspects of the work. Funders of the surveys that contributed data to the re-estimates were not involved in its design, in data analysis, data interpretation, manuscript preparation, or decisions on where, how or when to publish in the peer reviewed press. The study was approved by the Research Ethics Committee of the London School of Hygiene & Tropical Medicine (11208). The corresponding author had full access to all the data and had final responsibility for the decision to submit for publication.

## Results

Data from 1,355 districts in 31 countries were age- and sex-standardized and contribute an estimated 662 thousand cases (95% confidence interval (CI) 324 thousand-1.1 million) to the global total. Adjusting prevalence estimates by age and sex reduced raw district-level estimates by a mean factor of 0.45 (range 0.03-2.28). In countries where it was possible to adjust the datasets by age and sex, the country level backlog reduced by a mean factor of 0.56 (range 0.22-0.94) (Table 1). Estimates from PBPSs in 398 districts could not be standardized by age and sex because the original datasets were unavailable; the unadjusted backlogs in these districts totalled 781 thousand cases. We adjusted these estimates by multiplying them across the board by 0.45, yielding 411 thousand cases (95% CI 283-557 thousand). Based on new expert assessment of available data, it was determined that eight countries no longer have a trichiasis backlog. For six countries, local expert assessment generated revised, non-zero estimates. For one country, other published estimates[17] were used. Finally, there were seven countries for which the 2009 estimates were retained. Data included in the overall estimate had been collected from 2000-2016 (Figure 1).

**Figure 1.**
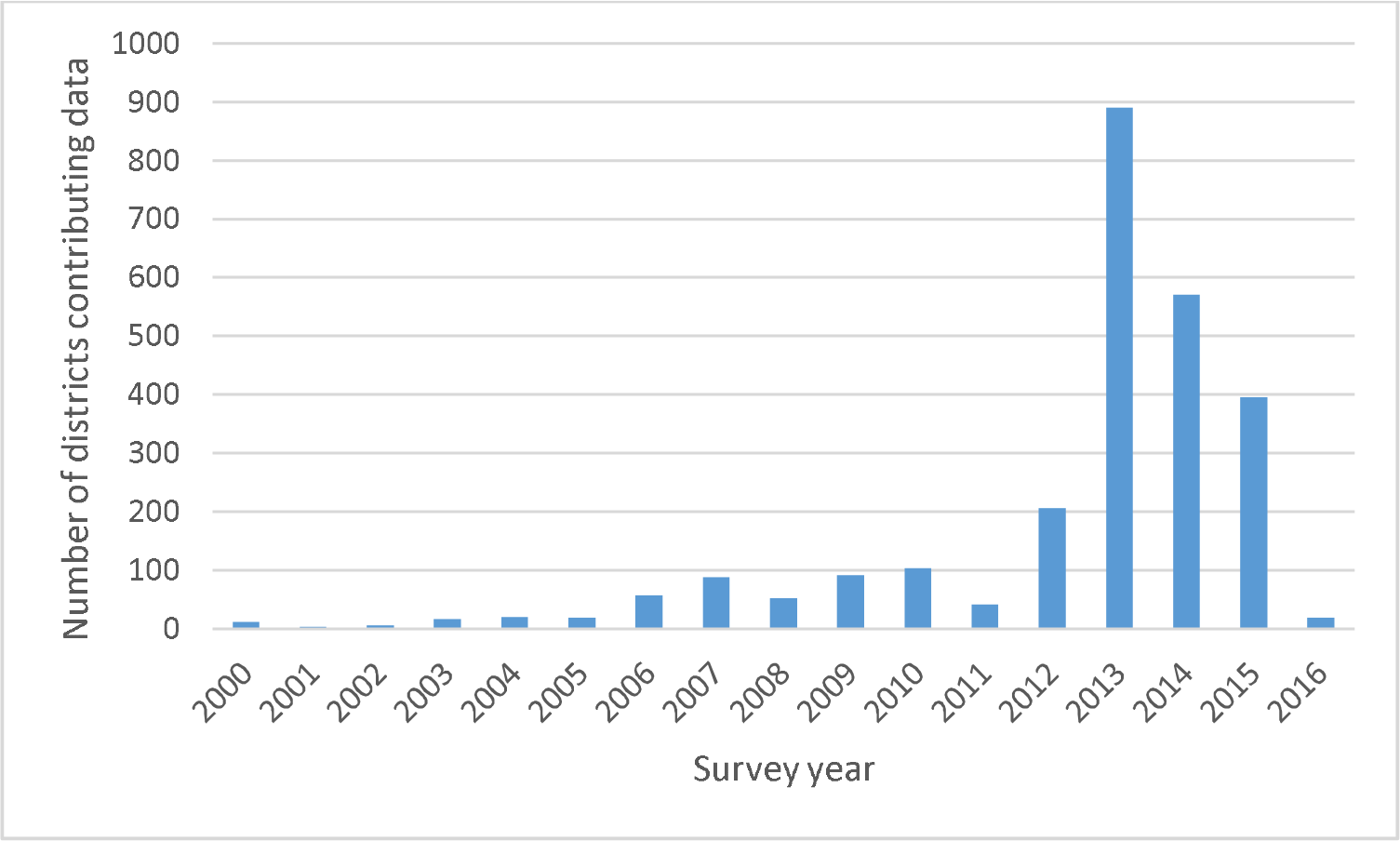
Number of districts contributing data to the overall estimate, by year of survey

**Table 1.** Estimated national-level trichiasis backlogs, 2016, with comparisons to the corresponding estimates for 2009

| WHO region | Country | 2009 estimate[8] (data generally from 2007) | Source for 2016 estimate (1) | 2016 estimate [year of data collection] (2) (95% confidence intervals) | Representativeness |
| --- | --- | --- | --- | --- | --- |
| African Region | Algeria | 86,700 | Retained previous estimate | 86,700 (23,022-175,625) |  |
|  | Angola | no data | Expert assessment (3) | 0 |  |
|  | Benin | 7,600 | GTMP[18] | 11,782 [2014-2015] (5,148-20,861) | Estimate based on 12 age- and sex-adjusted PBPS datasets covering all districts in the country where evidence indicates trachoma may be a public health problem |
|  | Botswana | 32,900 | Expert assessment (3) | 0 |  |
|  | Burkina Faso | 32,800 | PBPS – no raw data available | 19,443 [2005-2015] (5,381-35,236) | Estimate based on 63 PBPS datasets covering all districts in the country |
|  | Burundi | no data | Expert assessment (3) | 0 |  |
|  | Cameroon | 47,200 | PBPS – no raw data available | 698 [2012-2014] (27-1,523) | Estimate based on 1) 35 age- and sex-adjusted PBPS datasets, plus 2) 15 PBPS datasets, covering all districts in the country where evidence indicates trachoma may be a public health problem |
|  |  |  | PBPS – raw data available | 10,743 [2010-2014] (5,705-16,525) |  |
|  | Central African Republic | 1,000 | PBPS – no raw data available | 6,539 [2011] (1,991-11,259) | Estimate based on 8 PBPS datasets; security concerns prevented the remaining 7 districts in the country where evidence indicates trachoma may be a public health problem from being surveyed |
|  | Chad | 34,300 | GTMP | 24,597 [2014-2015] (8,513-44,924) | Estimate based on 1) 43 age- and sex-adjusted PBPS datasets, plus 2) 14 PBPS datasets. At the time of publication 1 district where evidence indicates trachoma may be a public health problem remained to be surveyed |
|  |  |  | PBPS – no raw data available | 23,024 [2002] (15,365-30,683) |  |
|  | Congo |  | GTMP | 0 [2015] | Estimate based on 1 age- and sex-adjusted PBPS dataset covering all districts in the country where evidence indicated that trachoma might have been a public health problem |
|  | Côte d'Ivoire | 59,900 | GTMP | 1,216 [2015] (176-2,582) | Estimate based on 10 age- and sex-adjusted PBPS datasets covering all districts in the country where evidence indicates trachoma may be a public health problem |
|  | Democratic Republic of the Congo | no data | GTMP[19] | 33,333 [2014-2016] (18,369-52,870) | Estimate based on 30 age- and sex-adjusted PBPS datasets;; security concerns prevented 1 district from being surveyed. At the time of publication an additional 9 districts where evidence indicates trachoma may be a public health problem remained to be surveyed. |
|  | Eritrea | 42,000 | GTMP | 774 [2014] (274-1,308) | Estimate based on 1) 2 age- and sex-adjusted PBPS datasets, plus 2) 36 PBPS datasets;; security concerns prevented the remaining 1 district in the country where evidence indicates trachoma may be a public health problem from being surveyed |
|  |  |  | PBPS – no raw data available | 24,793 [2006-2014] (15,005-34,823) |  |

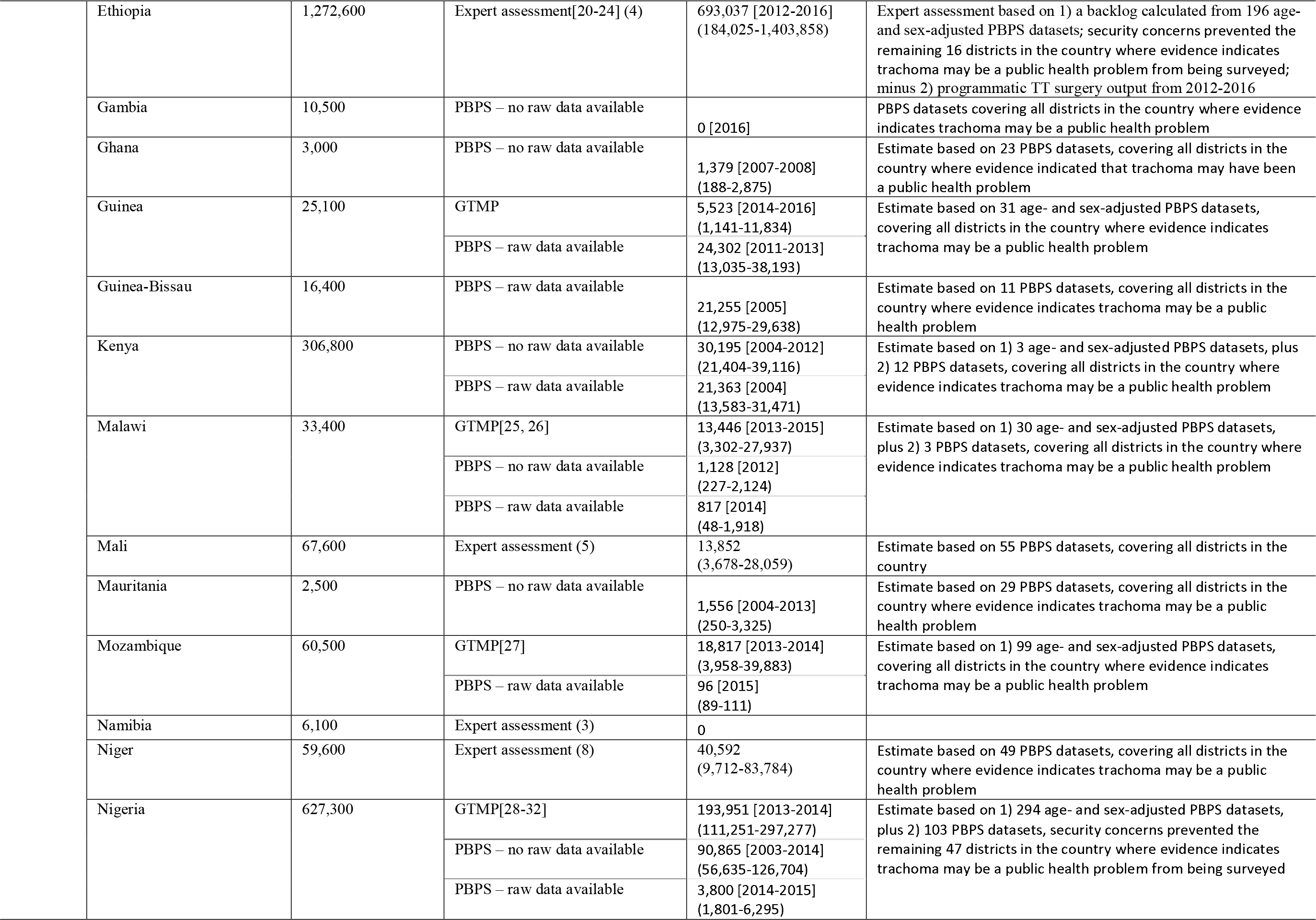

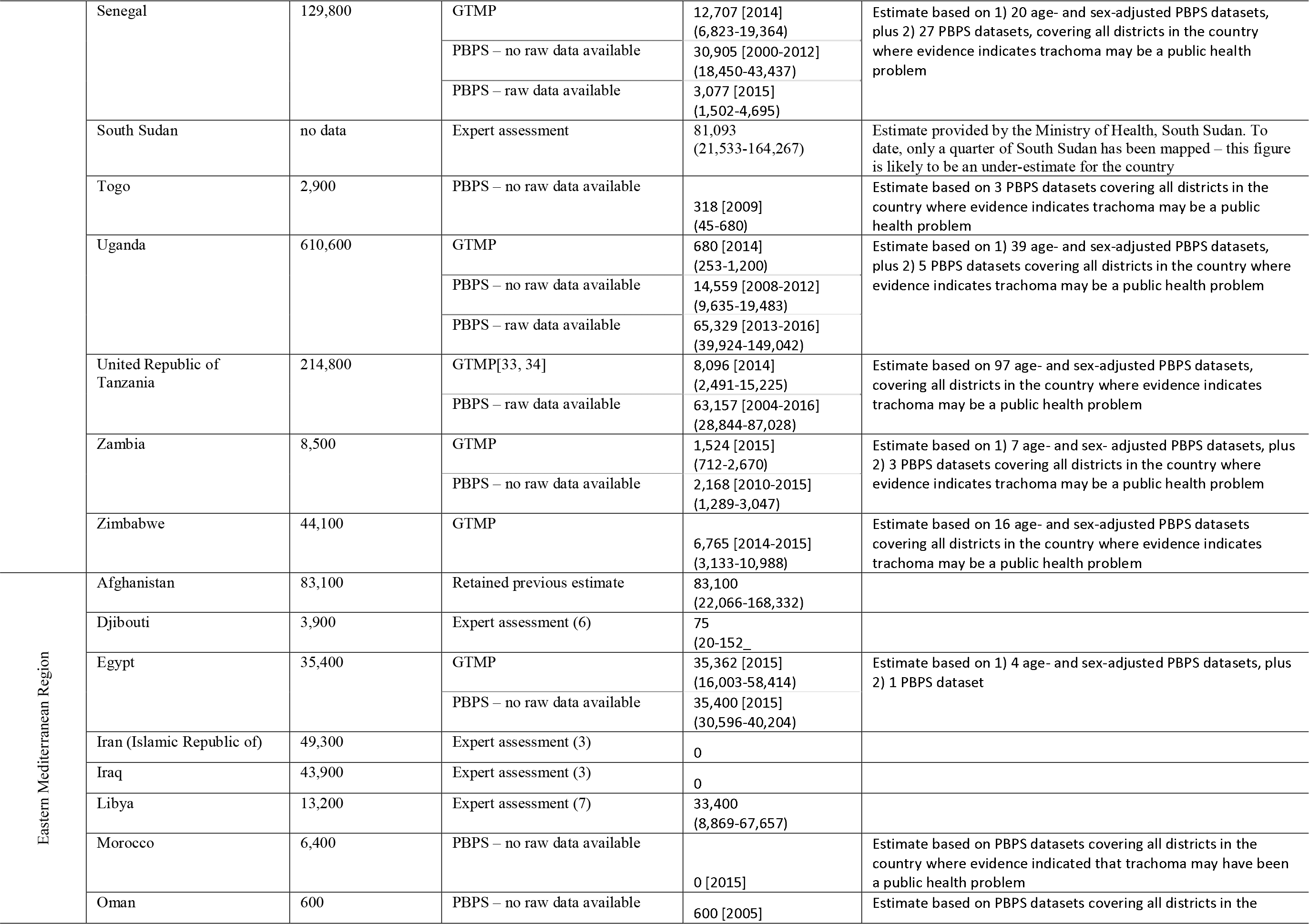

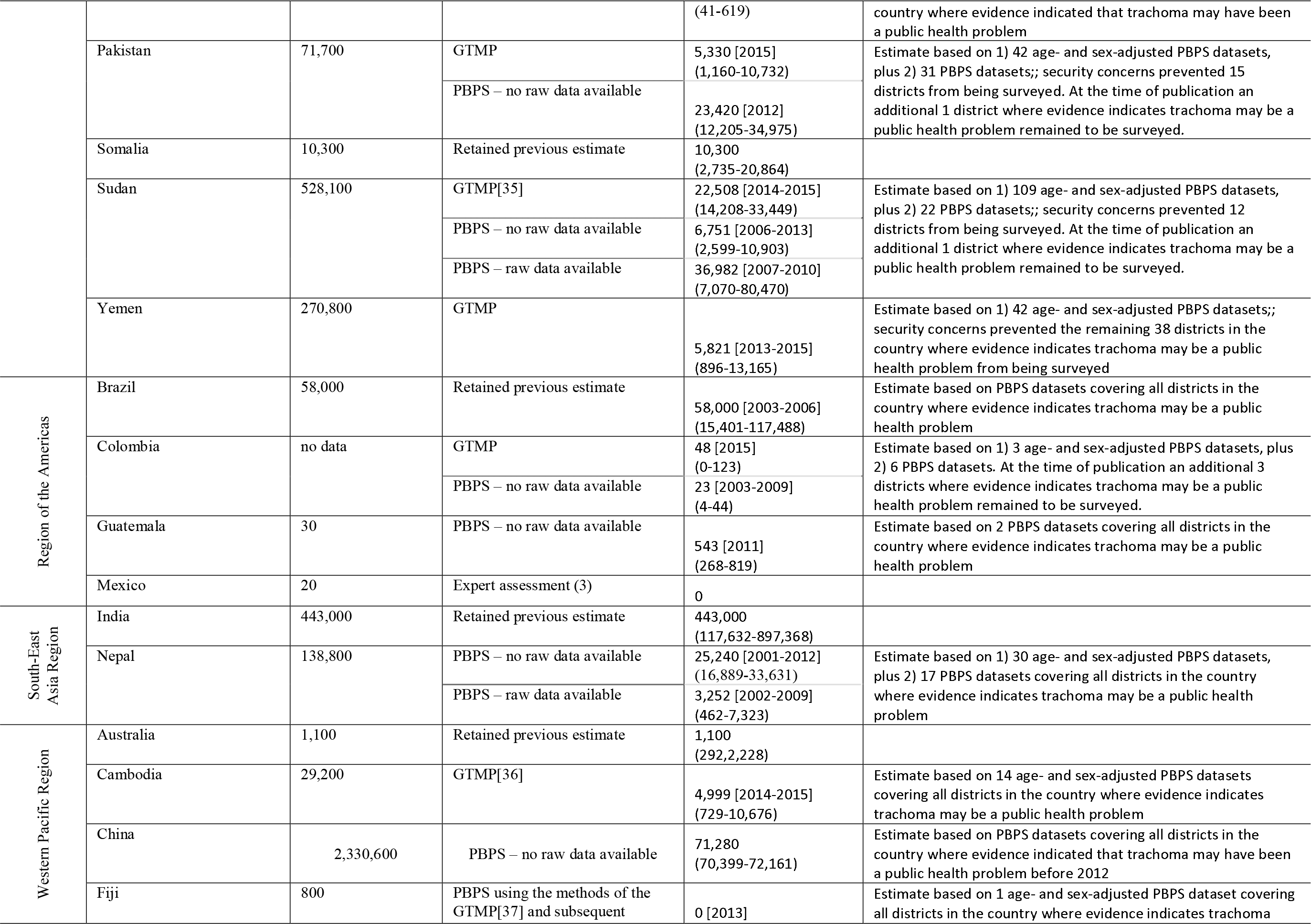

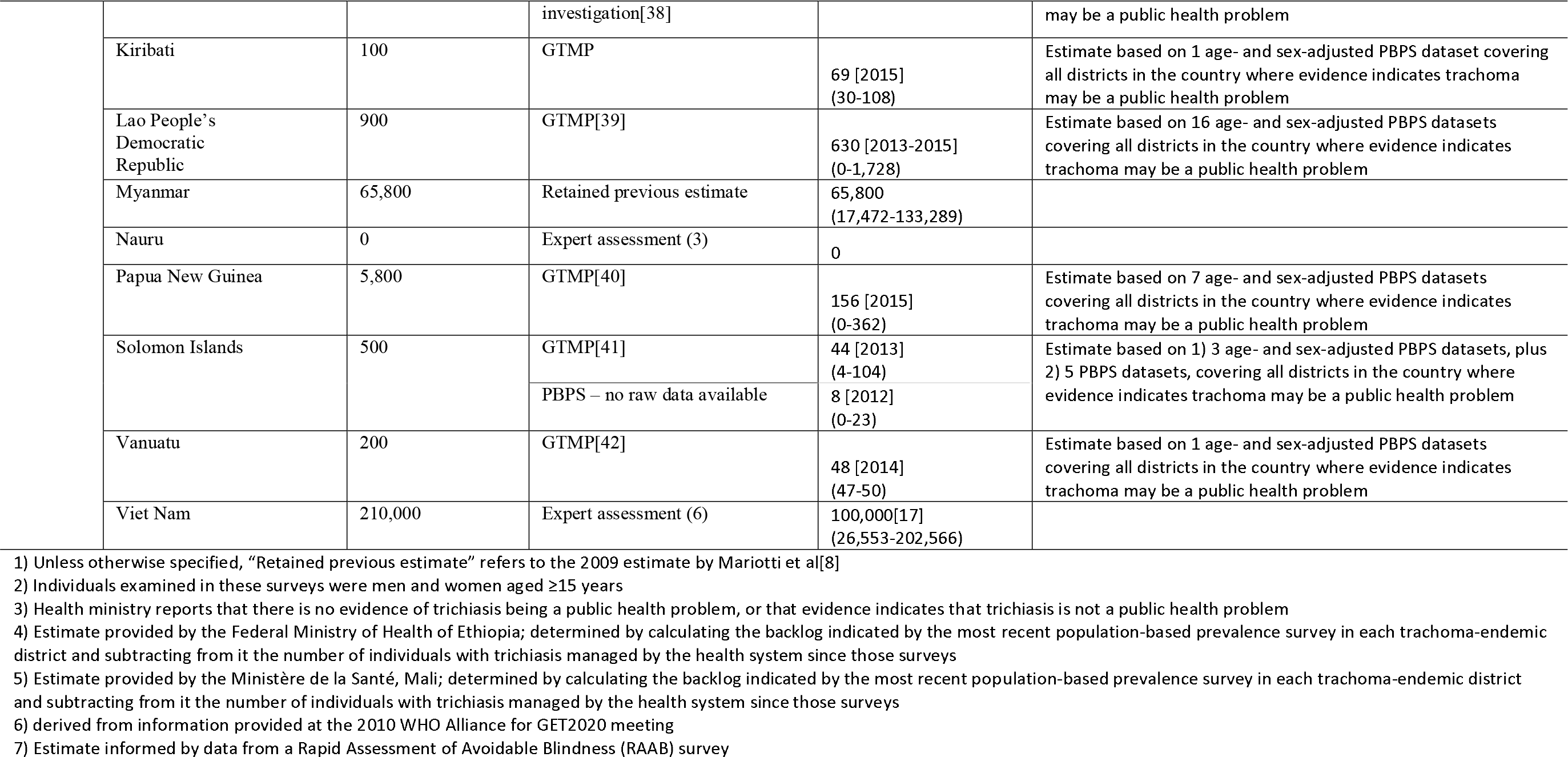

Through the inclusion of GTMP data, adjusting older datasets by age and sex, and obtaining current local expert assessment of available data, the global backlog estimate reduces from 8.2 million[8] (or 7.3 million[10]) to 2.8 million (95% CI 1.1-5.2 million) (Table 2).

In our estimates, data collected in surveys set up prior to August 2014 include all trichiasis, irrespective of the presence or absence of trachomatous conjunctival scarring. Data collected in surveys set up after August 2014 contributed data on trichiasis in eyes that also demonstrated trachomatous conjunctival scarring (or had an eyelid that could not be everted, due to presumed dense scar), thereby including only those cases of trichiasis attributable to trachoma. In both scenarios the data represent both “managed” and “unmanaged” trichiasis, irrespective of whether individuals have previously been offered corrective surgery or epilation[43].

**Table 2.** Estimated region-level trichiasis backlogs, 2016, with comparisons to the corresponding estimates for 2009

| WHO Region | 2009 Estimate | 2016 Estimate |  |  |  | Ratio (2016 estimate / 2009 estimate) |
| --- | --- | --- | --- | --- | --- | --- |
|  |  | Retained previous estimate or expert assessment (95% CI) | Unstandardized (reduced by a factor of 0.45) (95% CI) | Standardized (95% CI) | Total (95% CI) |  |
| African Region | 3,846,500 | 915,274<br>(241,969-1,855,593) | 247,570<br>(150,310-363,186) | 547,151<br>(283,052-913,841) | 1,709,995<br>(675,331-3,132,620) | 0.45 |
| Region of the Americas | 58,050 | 58,000<br>(15,401-117,488) | 566<br>(272-863) | 48<br>(0-123) | 58,614<br>(15,673-118,475) | 1.01 |
| Eastern Mediterranean Region | 660,000 | 126,875<br>(33,690-257,006) | 66,171<br>(45,441-86,702) | 106,003<br>(39,337-196,230) | 299,049<br>(118,467-539,937) | 0.52 |
| South-East Asia Region | 485,000 | 443,000<br>(117,632-897,368) | 25,240<br>(16,889-33,631) | 3,252<br>(462-7,323) | 471,492<br>(134,982-938,323) | 0.97 |
| Western Pacific Region | 2,630,000 | 166,900<br>(44,318-338,083) | 71,288<br>(70,399-72,184) | 5,947<br>(810-13,027) | 244,135<br>(115,527-423,294) | 0.09 |
| Total | 8,248,050 | 1,710,049<br>(453,010-3,465,538) | 410,835<br>(283,311-556,567) | 662,403<br>(323,661-1,130,543) | 2,783,285<br>(1,059,980-5,152,649) | 0.35 |

## Discussion

Trichiasis remains a significant public health problem in many countries, with a global backlog estimated for 2016 at 2.8 million people, 61% of whom live in sub-Saharan Africa. While there were methodological challenges in generating the 2009 estimate, it was the best estimate that could be made with the information then available. The considerable reduction from that estimate to the one generated here is likely to be the result of a combination of factors. First, there are now more and better data derived from rigorous surveys. Second, in many countries, there has been impressive programmatic scale up to manage TT, conducted by complex networks of governments and their partners. Third, there is likely to be an effect on the incidence of TT from the intensive efforts to reduce active trachoma prevalence in many contexts; such efforts have been ramping up in endemic countries since the World Health Assembly’s 1998 commitment to global elimination of trachoma[44]. Teasing out the relative contribution of each of these factors is not possible at the present time, but regardless of cause, the reduction is welcome news for global health.

A better understanding of the backlog and distribution of trichiasis cases is necessary to effectively plan for surgical services and other components of management of individuals with trichiasis. To reduce TT prevalence in each district of each endemic country to <0.2% in adults, which is the defined elimination threshold for TT[45], at least 2.0 million people will need to have their TT appropriately managed. From available data, it is estimated that 56% of people with trichiasis have bilateral trichiasis.

While our calculations lessen the uncertainty around the global backlog estimate, important limitations remain. Expert assessment of available data was used for 13 endemic countries, in seven of which the estimate was zero cases. Ethiopia, Mali, and Niger have undergone intensive surgical scale-up in the time since the most recent round of prevalence surveys. Because of this programmatic output, and a lack of consensus around how to counterbalance backlog reduction with new incident cases and post-surgical recurrence (both of which are inescapable, but presently difficult to accurately quantify) these countries provided results based on expert assessment. Uncertainty is greatest among the eight countries in which previous estimates were retained; these countries accounted for 848 thousand cases of trichiasis (30% of the global estimate) and further investigation is needed here. India, which accounts for 443,000 cases of trichiasis (52% of the total retained estimate) is a particular priority. A second ongoing uncertainty stems from the unavailability of raw data for 398 districts, for which, as a result, age- and sex-standardization was not possible. As noted in our calculations, there is a mean prevalence reduction against the raw prevalence estimate of ×0.45, which was applied to previously-unstandardized district-level estimates. Third, our CIs reflect statistical uncertainty. Because statistical uncertainty is uncorrelated across countries, our method of aggregating CIs across administrative boundaries overestimates statistical uncertainty at regional and global levels. However, there are other sources of uncertainty in addition to statistical uncertainty, so it is not possible to say whether our regional and global CIs are biased.

Ongoing lack of trichiasis (and TF) prevalence data for 235 suspected-endemic districts is due to local insecurity, and the global trachoma community stands ready to support national governments to undertake the needed mapping in those populations, when and if security conditions improve to allow safe conduct of fieldwork.

It is recognized[11] that the PBPS methodology described earlier is potentially imprecise in estimating TT against the WHO elimination target of ≤0.2% in adults. Thus, at district level, the uncertainty around the estimates of the TT backlog can be large. This is reflected in our CIs. Work is underway to try to develop more reliable methodologies for estimating the prevalence of trichiasis.

At present, the diagnosis of TT is based on the presence of clinical trichiasis (one or more lashes touching the globe or evidence of recent epilation of in-turned eyelashes) plus residence in a (presumed) trachoma-endemic setting. This definition is by nature somewhat circular and may need review, as it inevitably leads to classification of trichiasis as TT regardless of whether trachomatous conjunctival scarring is present or not. Trichiasis without trachomatous conjunctival scarring may be age-related or due to trauma, distichiasis, epiblepharon or other inflammatory disease[46]; the sight-threatening potential and optimal management strategies for non-trachomatous trichiasis still require further investigation[47].

Assumptions were made when adjusting the data for age and sex. First, we assumed that ages reported in surveys were accurate. However, it is reasonable to hypothesise that during a survey, individuals demonstrate a terminal digit preference when stating their age. Second, this analysis assumed that UNdata population pyramids used were representative of all districts for which survey data were available. Finally, because many studies have demonstrated a high correlation between trichiasis and increased age[48–51], we assumed zero trichiasis cases in the population aged 14 years and younger.

These calculations of national and global TT burdens are point-prevalence estimates based on data of differing vintages; they will change as additional surveys, baseline or impact, are undertaken. Other than for Ethiopia, Mali and Niger, the estimates do not take into account the number of surgeries done during the period between the most recent survey and this analysis. Nevertheless, estimates of current national TT backlog are essential for countries to appropriately allocate resources for surgical campaigns. Future district-level impact surveys in countries with trachoma elimination programmes will provide progressively improved evidence for countries to assess their residual TT burden and, ultimately, validate elimination of trichiasis as a public health problem. Baseline surveys of trachoma are still needed in a handful of countries (e.g., Egypt, Somalia and Central African Republic); these will lead to revisions in national estimates. In some settings, there will be a need to undertake trichiasis-only surveys in order to assess progress to elimination. Unfortunately, insecurity may continue to limit surveys in a few, but not all, of the settings in which previous estimates were retained. Where possible, surveys should be undertaken. Such data will help us drive towards, and demonstrate success in, the global elimination of trachoma as a public health problem.

